# The heart rate dynamics and antioxidant capacity under cognitive load in Obsessive-Compulsive disorder

**DOI:** 10.1101/2024.12.06.627156

**Authors:** Galina Portnova, Guzal Khayrulina, Elena Proskurnina, Rohit Verma, Olga Martynova

## Abstract

The research of the oxidants-antioxidants homeostasis and its connection to autonomic nervous system (ANS) dysfunction in patients with obsessive-compulsive disorder (OCD) is a relatively uncharted field of study. We recruited 31 patients diagnosed with OCD and 29 healthy volunteers to analyze their saliva antioxidant capacity before and after a three-block behavioral task with cognitive and emotional load, during which we assessed heart rate (HR) and heart rate variability (HRV) patterns. We found that the OCD group had a significantly lower antioxidant capacity (AOC) and showed no increase in AOC after the cognitive task as compared with healthy volunteers. The notable negative correlation between HR and AOC changes led us to select two subgroups inside the OCD and control group: one with high HR and another with low HR. The AOC changes for OCD and healthy participants with high HR were similar, while OCD sub-group with low HR exhibited significantly different changes in AOC compared with the control group. Although HRV did not show a correlation with AOC, the OCD group demonstrated substantially higher increase of very low frequencies (VLF) of HRV during the cognitive task performance and significantly elevated low frequency powers of HRV expressed in normal units (LFnu). OCD sub-group with lower HR exhibited higher LFnu compared to both the patient and the control group with high HR. Our findings imply that OCD patients with elevated sympathetic tone of ANS reflected by initially increased HR show more adaptive AOC to cognitive load

## Introduction

The etiology of OCD remains under investigation, and previous studies have suggested a variety of potential factors contributing to the development of this disorder [1]. These include genetic predisposition, experiences of emotional, physical, or sexual abuse or neglect, as well as social isolation, teasing, and bullying [1]. The symptoms of OCD could be easily revealed and affect the quality of life. Patients remain critical of their mental state, however, often develop an over-inflated sense of responsibility and magical thinking, an intolerance of uncertainty, and a belief in controllability [2] [3]. Chronic negative psychological influence accompanying the social activity of a patient with OCD can lead to life-long stress. Compulsion may be a maladaptive strategy for coping with a stressor that temporarily reduces the hyperactivation of the nervous system [4]. When the conditioned pattern of behavior has been fixed, it could be constantly repeated when obsessive thoughts arise. Since a person cannot get out of this vicious circle and this interferes with life and socialization, this subsequently aggravates stress. As a result, the chronic stress follows OCD and can be a trigger for the worsening of the OCD symptoms, accompanied by hypervigilance, increased negative affect, and mental arousal [5] affecting physiological reactions of the nervous and immune systems. The patient’s inability to cope with increasing pressure of chronic stress leads to an imbalance of sympathetic and parasympathetic tones of the autonomic nervous system (ANS) [6].

Utilizing electrocardiogram (ECG), altered values of the heart rate variability (HRV) and heart rate (HR) are frequently observed in mental disorders [7] and OCD is no exception. There are not many studies that reported HRV changes in OCD and current research findings are inconsistent or ambiguous. In particular, the severity of symptoms and the medication treatment could potentially lead to lower HRV of patients with OCD during resting state [8] [9]. At the same time, the studies, recruited patients with panic disorder, social phobia, and generalized anxiety disorder, did not show that the medications could affect HRV [9]. There were also conflicting findings regarding correlations between OCD symptom severity and HRV depending on whether HRV was measured in patients while sitting or standing [10] [11]. Recent study has revealed that patients with OCD showed higher HR and lower HRV compared to healthy participants [7]. Another study also reported that OCD patients were characterized by increased sympathetic and decreased parasympathetic tone [12]. Higher HRV in young adulthood distinguished individuals with OCD from healthy relatives, however lower HRV was indicative of relatives at older ages, according to an age-moderated group differentiation [7]. Previous findings in neurotypical teenagers suggested that low HRV, measured under respiratory sinus arrhythmia, acted as a moderator in the relationship between internalizing symptoms and psychosocial stress [13].

HR and HRV patterns may contribute to a more thorough understanding of the etiology of internalizing disorders like OCD [7] and could serve as a methodology to detect changes in the ANS parasympathetic and sympathetic activity in OCD [14]. The vagus nerve regulates HR centrally during rest, resulting in a reduction in HR [15] [16]. A low cardiac vagal tone may be a sign of internalizing psychopathological symptoms. HRV could be used to represent some features of parasympathetic regulation of the cardiac system [17]. This parameter has been also associated with higher levels of inhibitory control [18] [19] and voluntary emotion regulation [19] [20]. Since HRV is recovered following effective anxiety therapy, this proposes using HRV as an objective indicator of treatment response in OCD [14].

The associations between HR, emotional or cognitive arousal and antioxidant capacity (AOC) were previously reported in several studies. Some researchers showed an elevation of salivary antioxidant potential correlated with HRV and subjective pleasantness of stimuli after pleasant tactile stimulation in healthy participants [21]. The association between HRV and oxidative (malondialdehyde) responses were found in asthmatic children highlighting the inter-relationship between oxidative stress markers and ANS function [22]. Higher salivary antioxidant properties were also negatively correlated with HR in anthropometric studies of healthy children [23].

Oxidative stress may play a role in clinical manifestations of several psychiatric disorders including OCD and other anxiety and depressive disorders [24] [25] [26]. The importance of oxidative stress in the development of OCD could be explained by the dysfunction of catecholamines and the glutamatergic system in OCD and related disorders [27] [28]. Free radical damage to cells during the catecholamine metabolism process can result in abnormal neurotransmission at the dopaminergic neurons [29]. The studies investigated the oxidative stress or antioxidants in OCD demonstrated the importance of oxidative inflammation and lack of balance between the production of oxidative free radicals and neutralizing antioxidants in OCD and comorbid disorders [30] [31] hypothesizing the deficiencies of antioxidant enzymes in pure OCD [32]. Meta-analysis show that OCD patients have a systemic oxidative imbalance that is not adequately buffered by the antioxidant system [24].

In this study, we aimed to investigate the association between HR and HRV parameters and salivary oxidant-antioxidant activity in patients with OCD during a cognitive-emotional task. We hypothesized that, although there may not be significant differences in the baseline level of AOC between the control and OCD groups, the changes of oxidant-antioxidant activity after the cognitive load could vary and be associated with the baseline HR levels in OCD.

## Methods

### Participants

We used saliva samples and HR data of 29 healthy and 31 volunteers with OCD and 29 collected in the previous studies [4]. <u>The inclusion criteria</u> for the OCD group were the following: age from 18 to 40 years, right-handed, the diagnosis based on the criteria outlined in the International Statistical Classification of Diseases 10th Revision (ICD-10) (all participants were additionally interviewed by clinical psychologists (G.P., G.K.) to confirm their diagnosis); from 16 to 40 scores on the Yale-Brown Obsessive Compulsive Scale (Y-BOCS) – corresponding to moderate to extremely severe symptoms of OCD. <u>The exclusion criteria</u> for the OCD group were harmful/dependent use of substance, neurological or mental disorders other than OCD {bipolar disorder, autism spectrum disorder (ASD), schizophrenia}, traumatic brain injury, paroxysmal or epileptiform activity in electroencephalogram (EEG) and uncorrected vision. Patients with OCD were recruited from psychiatric clinics with a diagnosis of F42.0, F42.1 and F42.2 (ICD-10) for outpatient observation and treatment and were invited to a voluntary study in Institute of Higher Nervous Activity and Neurophysiology RAS (IHNA and NPh of RAS).

The control group of healthy participants was recruited via the announcement on social media. <u>The</u> <u>inclusion criteria</u> for control group were the following: age from 18 to 40 years, right-handed, participants were interviewed by a clinical psychologist to exclude any mental illness; from zero to 7 scores on Y-BOCS – corresponding to subclinical condition of OCD. <u>The exclusion criteria</u> for the control group were substance harmful use/dependence, neurological or mental disorders, uncorrected vision, abnormalities in EEG, or traumatic brain injury.

### Psychological assessment

All participants also completed the following questionnaires: Y-BOCS on web forms, State-Trait Anxiety Inventory (STAI), and Barrat Impulsiveness Scale (BIS-11).

### Experiment procedure

All participants were asked to have a good sleep the night before testing. Detailed instructions of the procedure were given to all participants. The cognitive and emotional load was simulated by a modified anti-saccade task with overlap design where affective pictures served as fixation stimuli [4]. The task consisted of three identical blocks lasting for 30-35 minutes. The whole experimental procedure with the preceding clinical interviews took 1.5-2 hours (Figure 1). The study was carried out using equipment of the Research Resource Center of IHNA and NPh RAS for functional brain mapping. The paradigm consisted of 300 trials of antisaccade task (100 trials per block). The stimuli and the paradigm were previously described in the study [4].

**Figure 1.**
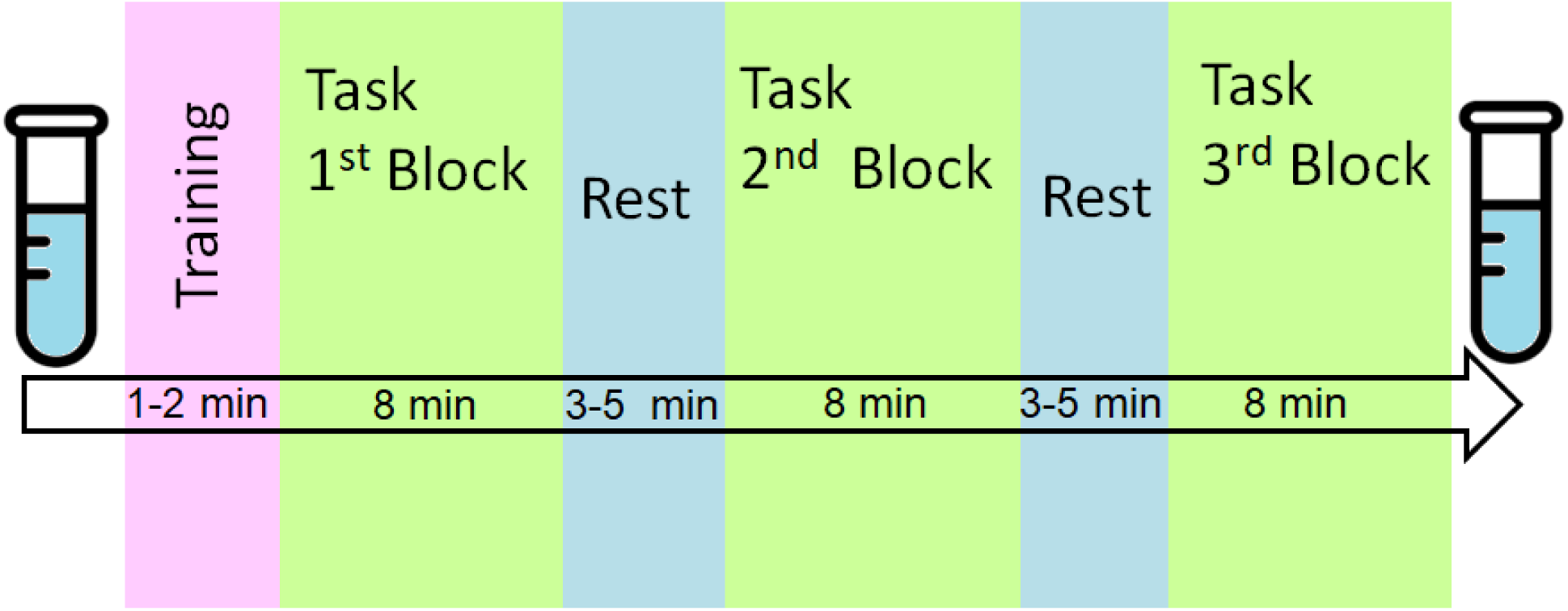
Experimental design

The volunteers were asked to take a comfortable seat in a chair, place a chin on a head mount and try to avoid large movements. Then calibration and validation of the eye-tracker camera were performed. The experiment was only started if the participant successfully passed the calibration and validation procedures (calibration was accepted if average error after validation was <0.30° of visual angle and maximum error was <0.50°). The task instruction appeared on the screen before every block. The participant could rest for about 3-5 minutes between blocks. Before proceeding with the next part, the participant passed the calibration and validation procedure again. The cognitive and emotional load while performing anti-saccade tasks was validated in our previous studies [4] [6].

### Saliva collection

The first saliva sample was obtained before the start of the experimental procedure; and the second saliva sample was obtained after the end of the experimental procedure (see Figure 1). The participants were warned about following specific rules to ensure a proper trace element analysis in saliva: to sleep at least 8 hours the day before the experiment, to stick to their usual daily routine, and do not take stimulants (caffeine immediately before the experiment, alcohol, drugs the night before) or other sedatives (if they are not related to the OCD medication therapy). Saliva was collected on an empty stomach by spitting. Before that, the participants rinsed their mouths with boiled water. The volume of saliva collected for the study amounted to at least 1 ml. Saliva samples were frozen at –20°C and stored for no more than a month before testing for trace elements.

#### Analysis of salivary antioxidant capacity

The AOC of saliva was quantified with the enhanced chemiluminescence protocol as described previously [33]. The chemiluminescent system consisted of a source of free radicals 2,2’-azobis (2-amidinopropane) dihydrochloride (ABAP, Sigma) and a chemiluminescent probe luminol (Sigma). A luminol solution of 1 mmol/L (Sigma) and ABAP solution of 50 mmol/L was prepared by dissolving the weighed samples in phosphate buffer solution (100 mM KH2PO4, pH 7.4, Sigma). The chemiluminescence was recorded at 37°C until a stationary level had been achieved, then an aliquot of 200 μL of saliva was added. The registration was performed until the new steady-state level was achieved. The antioxidant potential was calculated as the area of plot depression after the addition of the antioxidant to the system. The average of three parallel measurements was calculated. The measurements were carried out with a 12-channel Lum-1200 chemiluminometer (DISoft, Russia). Signal processing was performed with PowerGraph 3.3 Professional software.

### ECG acquisition and preprocessing

For all participants fifteen minutes resting-state ECG was recorded between 9:00 a.m. and 12:00 a.m. (before noon) or between 1:00 p.m. and 3:00 p.m. (afternoon) before the beginning of the cognitive task and ECG recording continued during 3 blocks of the active task solution. The recording cabin was temperature controlled (22–24 °C), dimly lit and the participants were comfortably seated. Bipolar ECG electrodes were placed at both wrists of the subjects; quality of data acquisition was visually tested before the start of the recording.

ECG data were recorded with the Brain Product amplifier (Gilching, Germany) using a blood pulse sensor attached at the middle finger of the right arm. The raw data after filtration 0.05 - 40 Hz was preprocessed with automatic R-peak detection using a Matlab script and a visual inspection of the identified R-peaks. Where necessary, an individual manual correction of R-peaks was applied. Datasets with artifacts in the time series were excluded from further analysis.

### 2.5. Heart rate patterns

Using the R-peak intervals, the time-domain, frequency-domain and non-linear parameters were calculated from the ECG data. The average HR was computed over three experimental blocks. HR was used to divide each group of participants into two subgroups according to their individual mean HR (low HR and high HR) as shown in Table 1. We used medians to divide the controls and OCD groups separately (Figure 4a). The median of the HR for the patients with OCD was 85.1, in low HR sub-group it was 74.6 and in high HR sub-group - 93.2 beats per minute. The average median of the in control group was 77.3, in low HR sub-group it was 70.0 and in high HR sub-group - 87.7 beats per minute.

**Table 1.** Inter and intragroup clinical and biochemical descriptive statistics.

|  | Patients with OCD (Valid N = 31) |  |  |  | Control group (Valid N = 29) |  |  |  |
| --- | --- | --- | --- | --- | --- | --- | --- | --- |
| Mean (SD) | All | High HR | Low HR | P value* | All | High HR | Low HR | P value* |
| Age | 23.1<br>(5.1) | 24.3<br>(4.8) | 22.9<br>(5.5) | 0.87 | 23.9 (6) | 23.5<br>(5.7) | 24.2<br>(5.3) | 0.99 |
| Y-BOCS<br>obsessions | 16.89<br>(4.6) | 16.88<br>(4.3) | 16.91<br>(4.7) | 0.99 | 6.14 | 6.53 | 5.69 | 0.68 |
| **Y-BOCS compulsions** | 11.04<br>(3.8) | 11.53<br>(3.5) | 10.27<br>(3.2) | 0.4 | 3.5 | 3.33 | 3.69 | 0.8 |
| **Y-BOCS Total** | 20.45<br>(4.4) | 22.28<br>(5.1) | 17.45<br>(4.1) | 0.16 | 1.18<br>(0.2) | 1 (0.1) | 1.38<br>(0.2) | 0.64 |
| **STAI** | 57.86<br>(17.4) | 57<br>(20.2) | 59.27<br>(16.8) | 0.41 | 45.36<br>(15.7) | 46.07<br>(15.1) | 44.54<br>(14.6) | 0.51 |
| **BIS-11** | 76.31<br>(21.1) | 77.44<br>(22.0) | 74.45<br>(20.4) | 0.31 | 66.29<br>(17.8) | 66 (19.1) | 66.62<br>(18.4) | 0.78 |
| **1<sup>st</sup> saliva sample AOC** | 29.07<br>(14.62) | 29.3<br>(11.2) | 26.8<br>(12.9) | 0.76 | 36.94<br>(14.73) | 41.4<br>(10.5) | 30.9<br>(11.9) | <0.01 |
| **2<sup>nd</sup> saliva sample AOC** | 30.92<br>(21.94) | 37.7<br>(15.2) | 20.9<br>(19.8) | <0.01 | 52.77<br>(15.29) | 52.2<br>(12.5) | 54.1<br>(11.7) | 0.69 |
| **HR** | 83.85<br>(10.17) | 92.29<br>(7.56) | 72.7<br>(6.97) | <0.01 | 78.69<br>(10.31) | 86.95<br>(7.56) | 69.24<br>(6.97) | <0.01 |

AOC: antioxidative capacity; BIS-11: Barrat Impulsiveness Scale; HR: Heart Rate; SD: Standard Deviation;
STAI: State-Trait Anxiety Inventory; Y-BOCS: Yale-Brown obsessive-compulsive Scale. The average values is indicated, standard deviation is given in parentheses: Mean (St.dev.).
\*Mann–Whitney U test between High HR and Low HR groups

For frequency domain markers, a Fast Fourier Transformation (FFT) was performed for assessing following variables [12]:

1) High-Frequency Power (HF): Fast Fourier Spectrum of frequencies 0.15–0.4 Hz that presents the activity of the parasympathetic ANS branch.
2) Low-Frequency Power (LF): Fast Fourier Spectrum of frequencies 0.04–0.15 Hz that present mainly the activity of the sympathetic branch and to some parts activity of the parasympathetic branch of ANS [34].
3) Very Low Frequency Power (VLF): Fast Fourier Spectrum of frequencies 0.0–0.04 Hz that present the activity of the parasympathetic ANS branch.
4) The normalized Low-Frequency Power (LFnu) was calculated as the percentage of LF from the sum of LF and HF of the R-peak time series. While LF reflects predominantly the sympathetic activity of the ANS, HF is a marker for the parasympathetic activity. The LFnu as the ratio of LF/(LF + HF) thus describes the proportion of the sympathetic activity of the total ANS activity and is complementary to the normalized High Frequency Power.

The following processing procedure was used to calculate these parameters. The sequence of intervals was converted into the corresponding beats per minutes (BPM) time series and then we used MATLAB’s spectrum function (Signal Processing Toolbox) to convert time series data to frequency data. The listed indicators 1-4 were calculated as the area under the resulting spectrum curve in the indicated ranges.

### 2.6. Statistics

All statistical analysis was performed using Statistica 13. The repeated measures mixed analysis of variance (rmANOVA) was used to evaluate effects of interaction between factor of group (2 levels) and within-subject factors of AOC (2 levels) and HRV dynamics (3 levels). rmANOVA was conducted for each of the HRV metrics. Post-hoc Bonferroni correction was employed. Furthermore, we conducted a correlation analysis (Spearman’s rank correlation coefficient) between HR, HRV, AOC metrics and BDI scores, YBOCS scores.

## Results

The **psychological assessment** results are presented in Table 1. The significant differences between OCD and control groups were found for the following questionnaires: 1) Y-BOCS scores (both obsessions and compulsions sub-scales) were higher in OCD group (Mann-Whitney rank test, r<0.45, p<0.001); 2) STAI scores were higher in OCD group (r=0.25, p=0.042). The differences between low and high HR sub-groups are depicted in Table 1 (see p-values). There were no significant gender or age differences between the groups.

### Salivary antioxidant capacity (AOC)

AOC was higher in healthy participants (group effect F(1, 58) = 15.997, p = 0.00018, ɳ2partial = 0.22). AOC increased during the study (F(1, 58) = 14.330, p = 0.00037, ɳ2partial = 0.20). However, this effect was significant only for the healthy volunteers (F(1, 58) = 8.9527, p = 0.00406, ɳ2partial = 0.13) as depicted in Figure 2.

**Figure 2.**
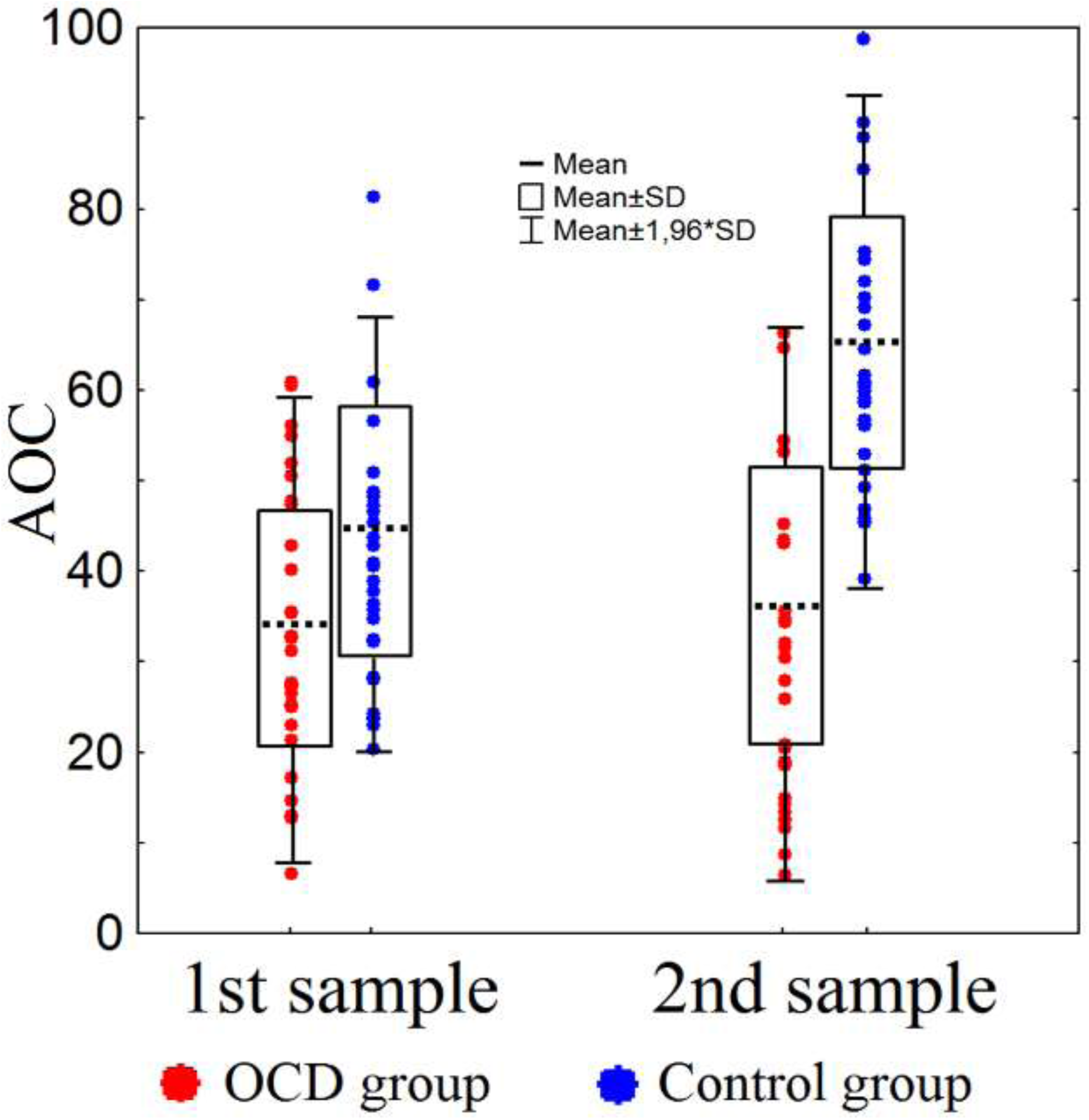
*The antioxidant capacity (AOC) dynamics in patients with OCD and healthy volunteers*

### Heart rate (HR) and heart rate variability (HRV) patterns

The mean HR was significantly higher in OCD group compared to healthy participants (group effect F(1, 56) = 4.8248, p = 0.03221, ɳ2partial = 0.08).

As depicted in Table 2, the VLF increased from first to second and third blocks of the task (main effect F(2, 116) = 73.522, p = 0.0000, ɳ2partial = 0.55). This effect was significantly more pronounced in the OCD group (mixed effect F(2, 116) = 3.892, p = 0.0242, ɳ2partial = 0.07): the VLF values in 2nd and 3rd experimental blocks were significantly higher in the OCD group (LSD test p < 0.0094).

**Table 2.** Heart rate variability parameters calculated separately in three blocks for each group of participants (OCD and Control group).

| group | Valid N | HRV parameters | Block1 |  | Block2 |  | Block3 |  |
| --- | --- | --- | --- | --- | --- | --- | --- | --- |
|  |  |  | Mean | SD | Mean | SD | Mean | SD |
| OCD | 31 | LF | 0.03 | 0.02 | 0.04 | 0.03 | 0.05 | 0.05 |
|  |  | HF | 0.02 | 0.02 | 0.03 | 0.03 | 0.03 | 0.03 |
|  |  | VLF | 23.92 | 9.96 | 34.94 | 12.23 | 36.93 | 27.82 |
|  |  | LFnu | 0.58 | 0.13 | 0.63 | 0.19 | 0.63 | 0.16 |
| Control | 29 | LF | 0.03 | 0.03 | 0.05 | 0.05 | 0.06 | 0.09 |
|  |  | HF | 0.02 | 0.02 | 0.03 | 0.02 | 0.03 | 0.03 |
|  |  | VLF | 23.17 | 6.73 | 31.89 | 9.71 | 31.86 | 10.34 |
|  |  | LFnu | 0.53 | 0.16 | 0.54 | 0.16 | 0.55 | 0.15 |

HF: High-Frequency Power; HRV: Heart rate variability; LF: Low-Frequency Power; LFnu: normalized Low-Frequency Power; SD: Standard Deviation; VLF: Very Low Frequency Power

Additionally, Friedman test followed by Wilcoxon Matched Pairs Test and Mann Whitney U Test was done: Friedman ANOVA Chi Sqr. (N = 83, df = 2) = 82.14 p<0.0001; Wilcoxon test between 1st and other blocks was z>7,18, p<0.0001; the 2nd and 3rd blocks didn’t differ (z =1.06, p=0.29); the VLF values in 2nd (z=2.00, p=0.045) and 3rd (z=2.14, p=0.03) experimental blocks were significantly higher in the OCD group.

The LFnu was significantly higher in the OCD group compared with the control group (group effect F(1, 58) = 8.2772, p = 0.00561, ɳ2partial = 0.12) as shown in Figure 3.

**Figure 3.**
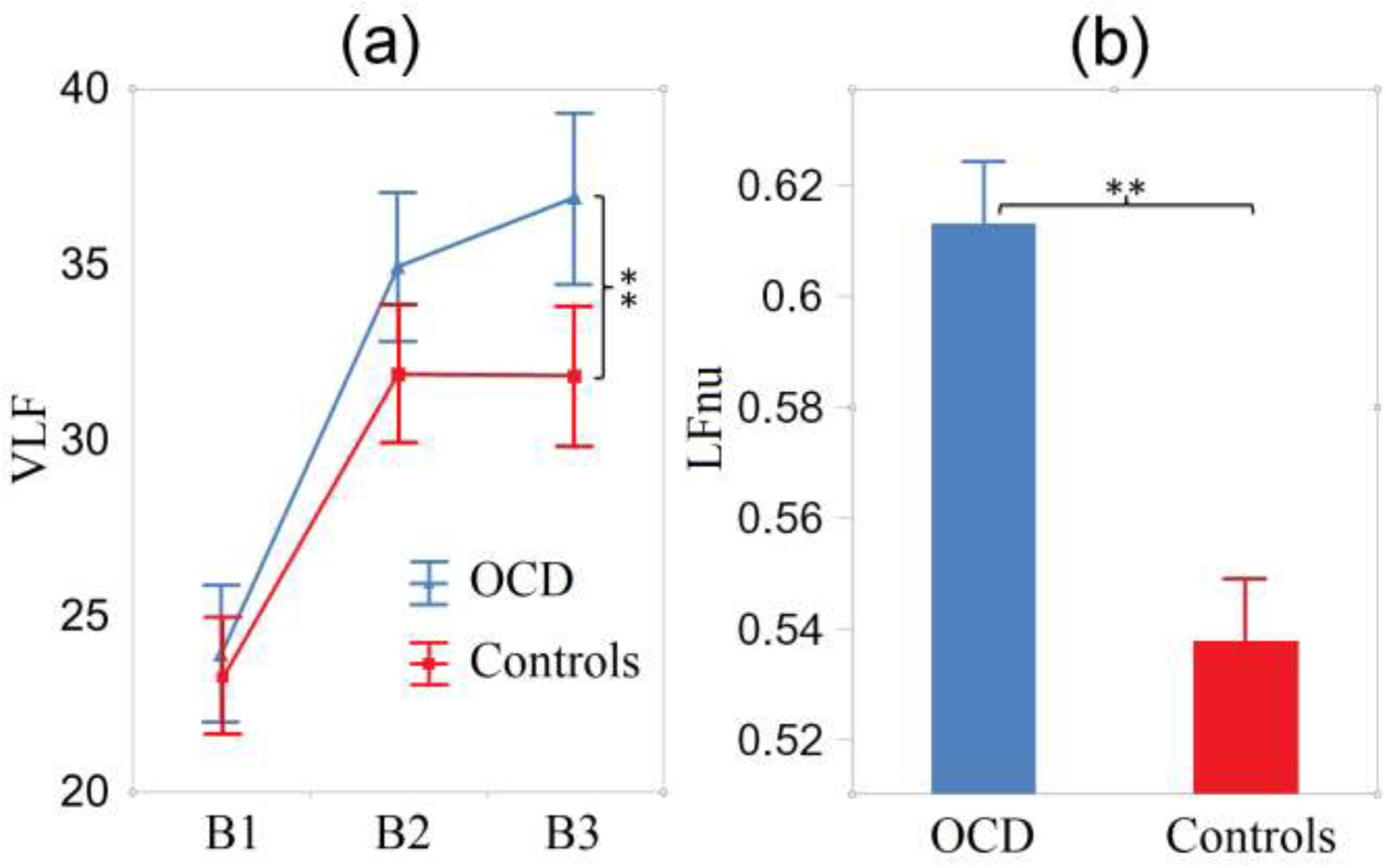
The HR data (a): the dynamics of VLF over three blocks in OCD and healthy volunteers; (b): LFnu in OCD and healthy volunteers.

### Correlation analysis

The correlation analysis prepared for the whole sample of participants (Spearman Rank Order Correlation) revealed significant correlations between questionnaires and AOC. The AOC in the 2nd saliva sample (r = -0.57, p < 0.0001) and the increase of AOC during the study negatively correlated with the scores of the STAI (r = -0.57, p < 0.0001 and r = -0.42, p = 0.001, correspondingly), scores of BDI (r = -0.57, p<0.0001 and r = -0.42, p = 0.0012, correspondingly), and scores of BIS-11 (r = -0.50, p < 0.0001 and r = -0.29, p = 0.0288, correspondingly).

The correlation analysis made separately in the OCD group and healthy participants showed significant association between HR and AOC. The mean HR positively correlated with AOC in the 2nd saliva sample (r=0.44, p=0.0029) and with the increase of AOC during study (r=0.38, p=0.016) in OCD. In the control group, the mean HR positively correlated with AOC in the 1st saliva sample (r=0.46, p=0.0016) and negatively correlated with the increase of AOC during the study (r=-0.58, p<0.0001).

We did not observe significant correlations of participant age with either AOC or HR patterns or questionnaires.

### Sub-groups with low and high baseline heart rate (HR)

The changes of AOC were oppositely related to the mean HR of participants (Wilks lambda = 0.85222, F(2, 55) = 4.7688, p = 0.01231, ɳ2partial = 0.15). The results are depicted in Figure 4(b). The high HR OCD sub-group (post-hoc: p = 0.0083) demonstrated a significant increase of AOC from the 1st to 2nd saliva sample, whereas individuals with OCD from the low HR sub-group demonstrated a trend of the decrease of AOC (p = 0.091). The healthy volunteers showed a significant increase of AOC from the 1st to 2nd saliva sample in both sub-groups, however in the lower HR control sub-group it was significantly more pronounced compared to the high HR control sub-group (post-hoc: p < 0.0001 and p = 0.0183, correspondingly). The average values of AOC (mixed effect F(2, 116) = 3.556, p = 0.03258, ɳ2partial = 0.08) differed between the OCD sub-groups (high HR and low HR) (post-hoc: p = 0.0079) but did not differ between control subgroups.

**Figure 4.**
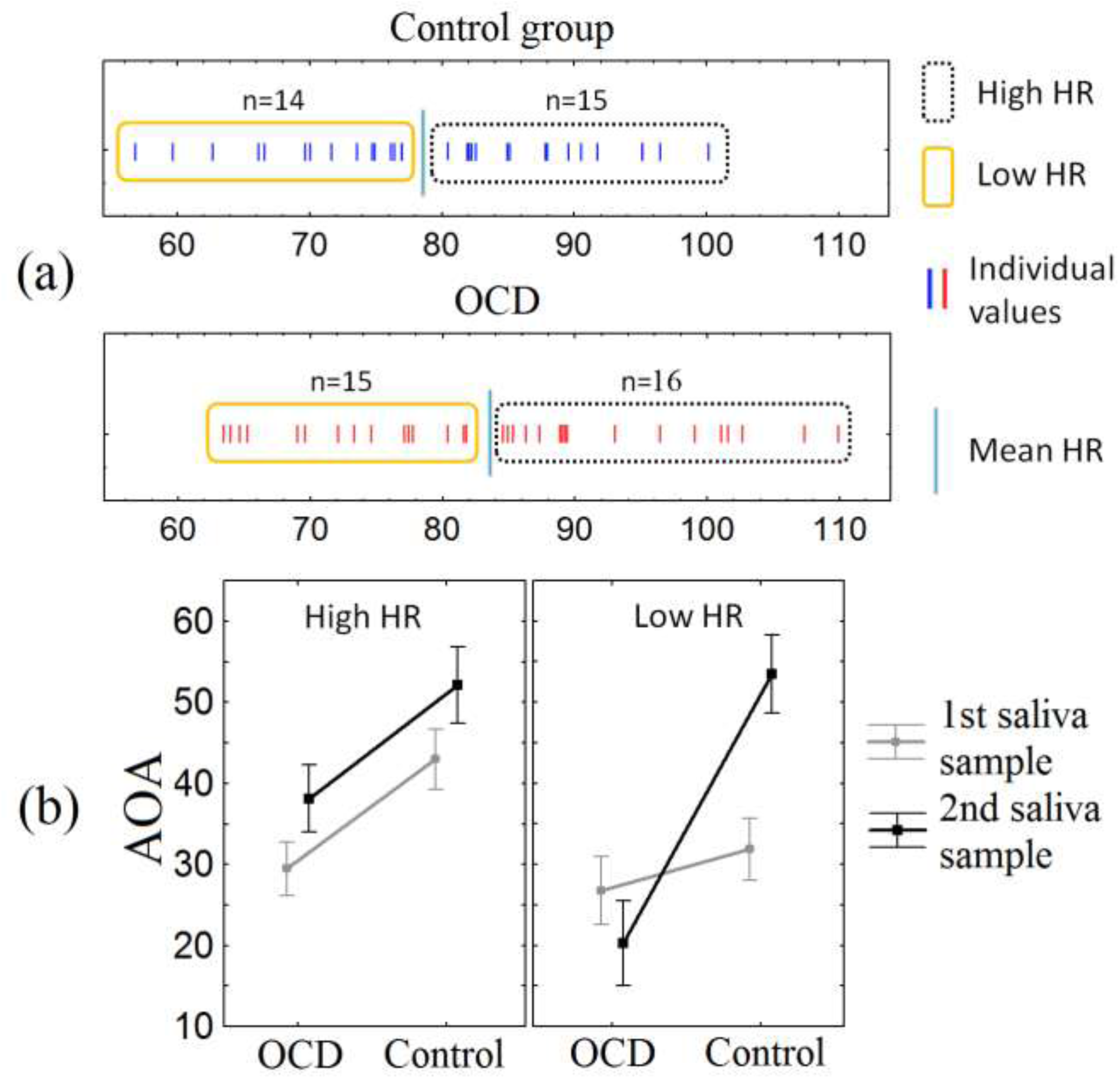
(a) The individual values of average HR and the scheme of sub-group division; (b) The changes of AOC in OCD and healthy participants with high and low HR

The higher and lower HR sub-group also differed in their LFnu dynamics during the study (see Figure 5). Individuals with OCD from the lower HR sub-group demonstrated a significant increase of LFnu in the 2nd and 3rd block compared to the 1st (post-hoc: p = 0.0067 and p = 0.0051), whereas in the other three subgroups the change of LFnu from block to block was not significant. Whereas the OCD sub-group with high HR had higher LFnu compared with the appropriate control group (F(2, 112) = 3.2408, p = 0.0429, ɳ2partial = 0.06) during all blocks of cognitive task performance, the OCD sub-group with low HR showed significant elevation of LFnu after the first block.

**Figure 5.**
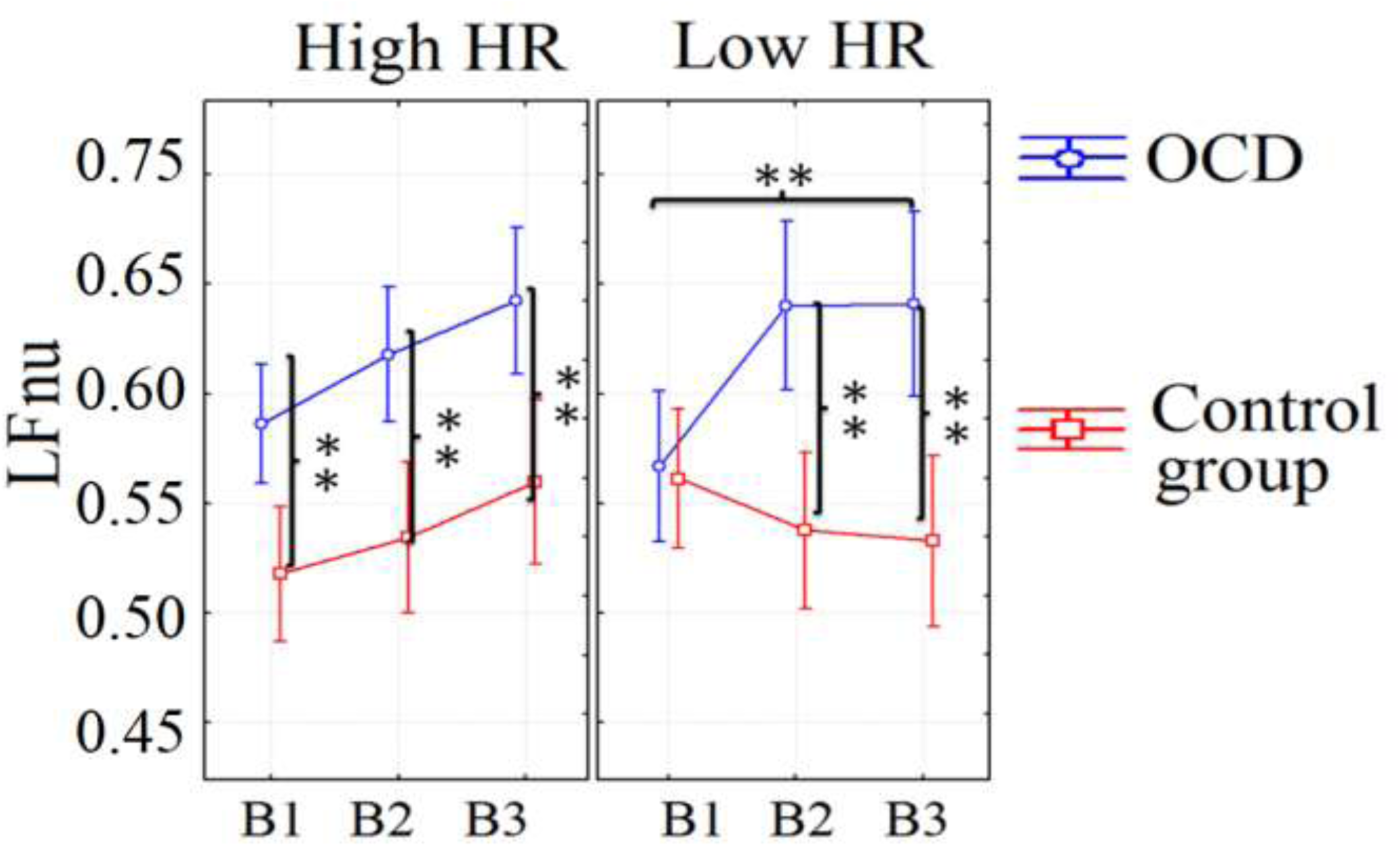
The LFnu dynamics in OCD and healthy participants with high and low HR

## Discussion

The AOC was significantly higher in healthy participants compared to the OCD group and increased significantly after the cognitive task performance also only in healthy volunteers. The imbalance of the oxidant-antioxidant activity and the antioxidant defense were previously shown to be associated with OCD [35]. Studies demonstrated the elevated level of products of lipid peroxidation, such as malondialdehyde in patients with major depressive disorder accompanied by OCD [32] and in patients with pure OCD [36]. Meta-analytic studies reported that patients with OCD demonstrated statistically significant increases in oxidant markers, but non-obvious results regarding antioxidant markers [24]; [37]. In contrary to our results, several studies found an oxidative imbalance shifted towards the antioxidant side and imbalance in antioxidant vitamins level in OCD [38] [36] [25]. At the same time, the use of antioxidants against oxidative stress is a novel treatment showed a significant improvement in the OCD symptoms [30].

Apparently, the relationship between antioxidants and OCD symptoms is complex and nonlinear, and depends on the compensatory abilities of the nervous system that are activated when coping with stress becomes necessary. Undoubtedly, one of the systems that regulate the normalization of stress levels is ANS. Our results demonstrated the opposite correlation between HR and the changes of AOC after solving the cognitive tasks in the OCD group and healthy volunteers which was positive in the OCD group and negative in controls. In the OCD group, the higher HR was associated with elevated levels of AOC after the task-solving period. Dividing the participants in sub-group with high and low baseline mean HR we additionally elucidated the opposite association between HR and antioxidant level in OCD compared with controls. Both sub-groups of the individuals with higher and lower HR from the control group demonstrated the increase of AOC after the task-solving; however, it was significantly more pronounced in participants with initially lower HR. Meanwhile, the OCD sub-groups with lower HR and higher HR differed significantly by the AOC changes and the average levels of AOC that were significantly diminished in the subgroup with lower HR. individuals with OCD with higher HR had the higher AOC level and demonstrated similar to the control group significant increase of AOC after the cognitive load. Our findings imply the higher adaptive functioning of ANS and oxidant-antioxidant system in OCD accompanied by elevated HR, who demonstrated ANS responses to be more similar to the controls as compared with OCD patients having initially low HR. According to the previous findings, HR in OCD could vary depending on different factors which are currently not entirely clear and, as a result, some studies reported the elevated HR in OCD [12] [7], while other studies did not find significant alterations of HR in OCD [9]; [11]. The significant correlation between elevated HR and the severity of OCD also was not observed. It is possible that the increased HR in patients with OCD is not necessarily a negative indicator; rather, it may serve as one of the compensatory mechanisms that encompass changes in both ANS and the oxidant-antioxidant system.

Regarding the HRV parameters, significant differences were found for the VLFnu and VLF patterns. The VLF component was previously associated with the inhibited parasympathetic activity or even the blockade of the parasympathetic activity of the ANS branch (Taylor et al., 1998). According to our results VLF increased in both groups during the study; however, the OCD group demonstrated a more pronounced elevation of this parameter. LFnu was also significantly higher in OCD compared with the control group. Although contemporary research prefers to view LFnu as a general indicator of aggregate modulation of both the sympathetic and parasympathetic branches of ANS, the majority of previous studies viewed this parameter as an index of modulation of the sympathetic branch of ANS [39].

Therefore, we assumed that the ANS activity could be characterized by the dominance of sympathetic tone, which increased to the end of the task-solving study and could even be accompanied by the blockade of the parasympathetic activity.

Our data are consistent with the previous findings showing that OCD is characterized by a decreased parasympathetic tone and abnormal sympathetic reactivity compared to controls (Sandhya, 2022; [12]. However, the previous findings were focused on dysfunction of the parasympathetic part of the ANS [10] [40]. In particular, some studies revealed the negative association between some parameters of HRV (including HF and RMSSD), which are considered as indicators of parasympathetic activity and the severity of OCD [10] [40]. The described relations between the lower activity of the parasympathetic branch of ANS and the severity of symptoms of OCD could be explained by the previously reported reduced serotonin transporter availability in OCD patients [40]; [41] [42]. While the research is still in progress, it is becoming clear that the effectiveness of certain medications must be considered along with a variety of other factors in the study of OCD including parameters of homeostasis, trace elements or hormones.

The functioning of the sympathetic and parasympathetic components of ANS is an interconnected process that should be viewed jointly when studying pathogenesis of OCD [42]. The different dynamics of HRV during our study in the OCD sub-groups with high HR and low HR could support this point of view. In the OCD sub-groups, the significant LFnu elevation after the first block was observed in case of the lowered HR, participants with OCD demonstrated the higher reactivity of this parameter that dramatically differed from the response of the control group with low HR. Thus, the dynamics of the ANS response to cognitive load was less balanced in OCD patients with lower HR, which may also indicate reduced compensatory abilities of ANS in this sub-group.

## Conclusions

We found that patients with OCD exhibited a significantly lower AOC compared to healthy volunteers after the behavioral study with cognitive load. Additionally, a significant increase in AOC after the cognitive task solving was observed only in the control group. A significant correlation between HR dynamics and AOC changes in response to cognitive loads prompted us to divide each group into two subgroups: those with high HR and those with low HR. Our analysis revealed that the AOC changes for OCD and healthy participants with high HR were comparable, while the OCD sub-group with low HR significantly differed in AOC as compared with healthy volunteers. Although HRV patterns did not correlate with AOC, OCD groups exhibited significantly higher reactivity to cognitive load in VLF and notably higher values of LFnu. Both parameters have previously been linked to the activity of the sympathetic branch of ANS. Moreover, OCD patients with lower HR demonstrated greater reactivity in LFnu compared to both the OCD and control groups with high HR. Our results showed that OCD patients with elevated sympathetic tone of ANS reflected by initially increased HR show more adaptive AOC to cognitive load

## Acknowledgements

We gratefully acknowledge Olga Kashevarova for technical assistance, Krystsina Liaukovich and Elizaveta Panfilova for help in collecting data. We are especially grateful to our volunteers who participated in the study.

## Author contributions

GP: Data analysis, Visualization, Writing – original draft, Writing – review & editing. G.K: Conceptualization, Data collection, Investigation, Visualization, Writing – review & editing. EP: Data analysis, Writing – review & editing. OM: Conceptualization, Funding acquisition, Methodology, Project administration, Resources, Supervision, Writing – review & editing; EV: Conceptualization, Writing – review & editing.

## Ethical statement

All experimental methods complied with the Helsinki Declaration’s guidelines. The study protocol (#0125022021) was approved by the Institute of Higher Nervous Activity and Neurophysiology’s Ethics Committee. Before the study, each participant signed a written consent form. Participants’ information was anonymous and private.

## Clinical trial number Not applicable Conflict of interest

The authors declare that the research was conducted in the absence of any commercial or financial relationships that could be construed as a potential conflict of interest.

## Data Availability Statement

Upon request from corresponding author

